# Identifying water stress response haplotypes in barley using latent environmental covariates

**DOI:** 10.64898/2026.05.04.722807

**Authors:** Zachary Aldiss, Stephanie M. Brunner, Bita Heidariask, Karine Chenu, Silvina Baraibar, Dini Ganesgalingam, David Moody, Shanice van Haeften, Lee Hickey, Yasmine Lam

## Abstract

**Purpose:** Genotype-by-environment (***G × E***) interactions represent a major obstacle to increasing genetic gain in crop breeding, with the underlying physiological drivers often remaining obscured within conventional statistical models. This case study presents a novel framework that transforms the latent factors from Factor Analytic (FA) multi-environment trial (MET) models into heritable quantitative traits, enabling the genetic dissection of adaptive response patterns.

**Methods:** A Factor Analytical Linear Mixed Model (FA-LMM) was fit to plot-level yield data for 1,036 barley genotypes across eight Australian trials.

**Results:** Correlation of the factor loadings with APSIM-simulated environmental covariates demonstrated that the second latent factor FA2 was strongly correlated with the Water Stress Index (***r* = −0.83**) during the critical flowering period, establishing water availability as the main biological axis of crossover ***G× E***. Genotypic scores for the derived traits, Overall Performance (OP) and Water Stress Response (WSR), were subjected to high-resolution haplotype-based mapping using local Genomic Estimated Breeding Values (GEBV).

**Conclusion:** This analysis successfully identified major genomic regions that accounted for a substantial proportion of the additive genetic variance. Gene Ontology enrichment of candidate genes within the top haploblocks implicated fundamental pathways related to energy homeostasis, root development, and stress response, with notable candidates including *FTsH11, BPS1*, and *TDP1*. The distribution of favourable Haplotypes of Interest (HOI) in elite cultivars suggested a historical signature of inadvertent selection for these adaptive mechanisms. This framework provides an explicit bridge between statistical modelling and functional genomics, offering breeders actionable genetic targets for accelerated development of climate-resilient cereals.

## 1 Introduction

The challenge of improving crop performance is frequently complicated by genotype-by-environment (*G* × *E*) interactions, which can reduce the efficacy of selection and limit genetic gain. While major adaptive traits such as phenology are effectively managed by breeders, a substantial portion of *G* × *E* remains a statistical ‘black box’. Conventional genomic selection (GS) models typically predict overall genetic merit, often without explicitly dissecting the biological mechanisms that confer resilience to specific environmental challenges. Consequently, a deeper and physiologically grounded understanding of the genetic basis for environmental adaptation is required to accelerate the development of climate-resilient cultivars.

To manage the complexity of *G* × *E* and spatial variation within a trial, factor analytic (FA) modelling approaches have become the standard for analysing multi-environment trial (MET) data (Smith et al. 2001). Treating genotype as a random effect, under the assumption that genotype effects are correlated across environments, is consistent with quantitative genetics where environments are regarded as traits (Hill and Mackay 2004). While an unstructured genetic variance matrix can model every parameter, FA models offer a more parsimonious approach that capture the common genetic patterns across the MET datasets, with environment-specific variances accounting for the remaining lack of fit (Smith et al. 2001). Over the past two decades, selection accuracy has improved by leveraging FA modelling, consistently outperforming other modelling approaches (Kelly et al. 2007; Smith et al. 2015). However, these latent factors, which capture underlying and unobserved variables which influence the observed data, are typically used to enhance predictive accuracy, while their potential to provide direct biological insight remains largely untapped. Recent efforts to interpret these factors have relied on post-hoc analysis, such as the estimation of Pearson’s correlations between environmental loadings and environmental covariates averaged across the growing season to identify probable drivers of *G* × *E* (Oliveira et al. 2020).

The latent factors from an FA model are biologically driven selection indices. The first factor (FA1) typically represents the main genetic effect on performance or yield potential across the MET serving as an index of overall adaptation (Smith and Cullis 2018). Factors following FA1 capture the primary axes of *G*×*E*, serving as quantitative indices of specific adaptive responses that affect genotypic re-ranking across environments (Smith et al. 2015). Genotypic scores on these factors quantify how strongly a particular genotype expresses that specific adaptive response (Bančič et al. 2023). This provides a powerful opportunity to move beyond the analysis of yield in individual environments and toward dissecting the genetic control of these fundamental response patterns.

While FA models statistically describe these adaptive responses, the identification of underlying genetic drivers remains the primary objective. *G* × *E* interactions are fundamentally caused by loci with differential effects across environments, leading to the re-ranking of genotypes and conferring local adaptation (Marschner and Schou 2019; Via and Lande 1985). The challenge has been to connect the statistical patterns revealed by FA models to these causal loci. Similar approaches have recently integrated the environmental dimension into genome wide associate studies and genomic selection by defining reaction-norm parameters along a performance-free environmental index, which facilitates the forecasting of performance in new environments (Li et al. 2021). However, such methods often rely on population means which may obscure individual genotype sensitivities to latent environmental drivers. The central novelty of this research involved treating genotypic scores on each latent factor as novel and heritable quantitative traits. This establishment of these *G* × *E* interactions as heritable quantitative traits provides a critical bridge for applying powerful genomic mapping tools to dissect the genetic architecture of adaptation itself.

By defining these responses as traits, haplotype-based mapping can be leveraged to identify the chromosomal segments that control them. Haplotypes can be defined in many ways, though the most common method utilizes linkage disequilibrium (LD) or physical proximity on the chromosome (Voss-Fels et al. 2019). By analysing haplotypes rather than individual SNPs, haplotype-based approaches capture the effects of multiple linked loci that contribute to *G*×*E* as well as account for the effects of epistasis and gene-gene interaction. Using a local genomic estimated breeding value (GEBV) approach, genetic merit conferred by each haploblock can be estimated for these novel traits (Shaffer et al. 2025). This provides a high-resolution map of the genetic architecture controlling yield potential and specific environmental responses, allowing breeders to target superior haplotypes for selection within their target population of environments (TPE).

This framework was operationalised by combining a Factor Analytical Linear Mixed Model (FA-LMM) with APSIM crop modelling simulations. Biological meaning was first assigned to the primary statistical drivers of G×E in a large-scale barley MET dataset. Novel quantitative response traits which correlated to yield potential and plasticity to water stress were then derived using bootstrap correlations against simulated indices, such as the Water Stress Index (WSI) centered on the precise flowering window. Finally, the genetic architecture of these response traits was dissected through haplotype mapping to identify the underlying specific genomic regions and candidate genes, providing a clear path from complex field data to actionable breeding targets.

## 2 Materials and Methods

### 2.1 Plant materials and data sources

This study evaluated a panel of 1,035 barley genotypes, including both diverse and elite breeding material representing over fifteen years of commercial breeding history. The genotype and phenotype data were provided by InterGrain Pty Ltd. This panel was evaluated across eight rainfed environments (defined as a combination of location and year) in Western Australia and Queensland during the 2021, 2022 and 2024 growing seasons (table 1). A randomized block design was used within each trial, incorporating row and column blocking to account for spatial heterogeneity. Each trial had a mean of 633 unique genotypes in total with 480 genotyped. All trials were fully replicated except GAT24 with approximately 78% replication (table S1). There was a high level of genotype concurrence across all trials, with an average of 64% (table S2).

**Table 1.** Summary of field trial information.

| **Trial** | **Location** | **Lat.** | **Long.** | **Sowing** | **Harvest** | **Flowering** |
| --- | --- | --- | --- | --- | --- | --- |
| **CGN21** | Corrigin, WA | −32.3682 | 117.9402 | 22/05/21 | 17/11/21 | 07/09/21 |
| **CGN22** | Corrigin, WA | −32.3800 | 117.9448 | 26/05/22 | 23/11/22 | 16/09/22 |
| **YRK21** | York, WA | −32.0398 | 116.8339 | 17/05/21 | 30/11/21 | 28/08/21 |
| **YRK22** | York, WA | −32.0254 | 116.8626 | 19/05/22 | 15/11/22 | 24/08/22 |
| **DND21** | Dandaragan, WA | −30.7519 | 115.6683 | 12/05/21 | 16/11/21 | 16/08/21 |
| **DND22** | Dandaragan, WA | −30.6515 | 115.7843 | 05/05/22 | 14/11/22 | 05/08/22 |
| **GAT22** | Gatton, QLD | −27.5444 | 152.3578 | 16/06/22 | 10/11/22 | 04/09/22 |
| **GAT24** | Gatton, QLD | −27.5482 | 152.3269 | 21/05/24 | 25/10/24 | 21/08/24 |

Grain yield was recorded at recorded at maturity for each plot using plot harvesters with yields expressed as tonnes per hectare (t/ha).

Of the 1,035 accessions, 648 were genotyped using the Illumina Infinium 40K XT SNP chip. After quality control, which included removing markers with > 10% missing data and a minor allele frequency of < 0.01, a high-quality set of 6,733 polymorphic single nucleotide polymorphisms (SNPs) were curated for downstream analysis (figure S1).

### 2.2 Determination of environmental covariates

In order to quantify commonalities in environmental stressors between the environment experienced by the crops in each trial, simulations were performed for the commercial cultivars RGT Planet and Spartacus CL and characterised with the APSIM-Barley (version 2024.6.7514.0) crop model (Holzworth et al. 2018, 2014). Daily weather data from the nearest meteorological station in the SILO based database (https://www.longpaddock.qld.gov.au/silo/) (Jeffrey et al. 2001), data from nearby characterised soil in the APSoil database (http://www.apsim.info/Products/APSoil.aspx) and trial-specific management practices were collected. For Gatton 2022 and 2024 trials, initial soil moisture at different depths was measured onsite at sowing, and simulations commenced from the sowing date. For environments without measured soil moisture information (Corrigin, York, and Dandaragan), simulations began six months prior to sowing, with an initial plant available water of 20% uniformly distributed through the soil profile. To best simulate crop growth and development at each trial, an APSIM parameter of the cultivars (‘minLN’, i.e. the number of leaves that the crop produces under long photoperiod when vernalisation requirements have been satisfied) was tuned so that simulated flowering time matched observed flowering time (Amin et al. 2025).

A water stress index (WSI), defined as the ratio of crop water status supply to demand was simulated daily for the eight InterGrain trials (Chenu et al. 2013). Average temperature and WSI were calculated for 200 degree-days brackets centred at the simulated flowering time.

### 2.3 Statistical analysis

A single-stage, Factor Analytic Linear Mixed Model (FA-LMM) was applied to the raw plot-level yield data. A genomic relationship matrix was utilised and was scaled and centered according to VanRaden (2008) to account for non-full rank matrices. This approach was selected because it is an industry-standard practice for MET analysis, as it simultaneously models the additive genetic main effects and the complex genotype-by-environment (G×E) interactions, thereby improving the accuracy of genetic predictions in breeding programs (Smith et al. 2004). The model was fitted using ASReml-R v4 (Butler et al. 2017). The most optimal model was determined by evaluating the Wald test statistics for fixed effects, the highest Restricted Maximum Likelihood (REML) log-likelihood value, and the smallest Akaike Information Criterion (AIC). A FA-LMM of order three was used based on model fit statistics and additive genetic variance explained averaging 93.5% across all sites (table S4) and 58.26% of the total genetic variance (table S3). The general form of the model for the yield phenotype, y, was defined as

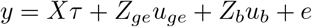

Where is the fixed effect term of each trial (environment), fitting a unique mean for yield for each location x year combination. Random blocking effects *u*_*b*_ contain row and column independently fit for each trial, with *u*_*b*_ ∼ *N* (0, *G*_*b*_). The vectors of random effects, uge containing the genotype by environment effects where a separable variance structure was assumed for *G* × *E* effects *u*_*ge*_ ∼ *N* (0, *G*_*e*_*G*_*g*_) where *G*_*e*_ is the variance-covariance for the environmental effects and *G*_*g*_ is the variance-covariance matrix for the genotypic effects. *G*_*g*_ is partitioned into additive and non-additive genetic variance matrices using genomic information for the 668 genotypes where SNP information was available. For the additive *u*_*ge*_ a factor analytic model of order three (FA(3)) was fit following the form *G*_*ea*_ = ΛΛ^⊤^ to account for the environment loadings and specific variances respectively. For the non-additive component an identity matrix was used for all genotypes and was fit to a factor analytical model of order one (FA(1)). The residual error, *e* was specified with a trial-specific, separable first order autoregressive (AR1*AR1) structure for column and row to model local spatial variation. The variance matrices for blocking effects *G*_*b*_(*σ*_*b*_) is specified as direct sums of the scaled identity matrix.

### 2.4 Latent regression

To identify the environmental drivers of *G* × *E*, the interaction was modelled using a FA-LMM framework. This approach effectively fits a latent regression model where the *G* × *E* interaction effect 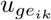 for genotype *i* in environment *j* was defined as:

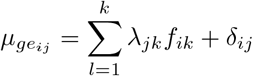

where *λ*_*jk*_ is the latent environmental covariate (loading) for the *k*-th factor in environment *j, f*_*ik*_ denotes the genetic sensitivity (score) and *δ*_*ij*_ represents the residual *G* × *E* variance specific to environment *j*. The genetic variance-covariance matrix across environments (*C*_*e*_) was derived from the FA parameters and decomposed as:

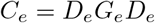

where *D*_*e*_ is the diagonal of the genetic variance within environments and *G*_*e*_ denotes the genetic correlation matrix between environments. The elements of Ge provided insight into the nature of the interaction. Positive correlations approaching unity indicated scale inter-actions (non-crossover), whereas negative correlations indicated rank changes between environments (crossover interaction). To allow interpretability, the factor solution underwent varimax rotation. This singular value decomposition applied to the estimated site loadings orthogonalised the factors while ensuring the first factor captures the maximum portion of genetic variance, with the second accounting for the remaining and so on. Latent regression plots were generated for each factor by plotting the Best Linear Unbiased Predictions (BLUPs) of the total genetic effects against the rotated latent environmental covariates (*λ*). The slope of this regression corresponded to the rotated genetic score for each genotype. These rotated scores were extracted for downstream analysis and haplotype mapping. To validate the biological basis of the latent factors, the environmental loadings were correlated with explicit environmental covariates. These explicit covariates were defined as external physical parameters, specifically cumulative rainfall, temperature stress, and radiation, which were simulated for each environment using the Agricultural Production Systems sIMulator (APSIM) model at discrete crop growth stages. An empirical permutation test was employed to determine significance of the observed correlations. For each covariate, the site level data was randomly permuted one thousand times to construct a null distribution with random gaussian equal to five percent of the standard deviation injected for each permutation. Empirical p-values were derived by calculating the proportion of the null correlations that were greater than the absolute magnitude of the observed correlation.

### 2.5 Haplotype mapping

To identify chromosomal regions associated with the genotypic sensitivity scores (*f*_*ik*_) and site-specific BLUPs, a haplotype-based approach was used to calculate local Genomic Estimated Breeding Values (local GEBV) (Shaffer et al. 2025). Prior to analysis, genome-wide SNP markers were grouped into haploblocks of high linkage disequilibrium (LD) based on a pairwise *r*^2^ value greater than or equal to a minimum threshold of 0.4. A marker tolerance of *t* = 3 was set per block to accommodate incorrectly positioned markers. A block was considered complete when at least three flanking markers fell below the *r*^2^ threshold. This process resulted in the formation of 1,145 haploblocks with an average of five markers per block (Supplementary Data 1).

Global marker effects for the response traits were predicted using the ridge regression linear unbiased prediction (rrBLUP) model (Endelman 2011). The local genomic estimate was derived by summing the marker effects within each haploblock to capture the cumulative additive effect of the haplotype. The variance for each haploblock was calculated as the range in haplotype effects for each haploblock. High impact regions or ‘top blocks’ were identified as those residing within the top 0.5% of the genome wide haploblock variance distribution. The top 0.5% of high-variance blocks were selected for further analysis, and their physical positions were cross-referenced against known quantitative trait loci (QTL) and genes related to drought adaptation and other relevant traits. A linkage map was generated using MapChart.

Haploblock variances were plotted in the format of a Manhattan plot to visualise the most important blocks in the barley genome. The top 2 to 3 blocks per analysis were also explored for their contribution to genetic variance by fitting a modified version of the FA3 model with a marker matrix composed solely of markers present in the relevant blocks. The genetic variance accounted for by this matrix was then contrasted with the genetic variance accounted for by the full GRM.

To translate these findings to the cultivar level, specific lines were screened for Haplotypes of Interest (HOIs). HOIs were defined as allelic combinations that exhibited a positive scaled additive effect on the target trait and maintained a minor allele frequency (MAF) ≥ 0.05 within the breeding panel. Given that haplotype effect distributions typically exhibited bimodal or multi-modal patterns, HOIs were identified by selecting the allelic clusters corresponding to the most positive peak in the density distribution.

### 2.6 Haploblock specific gene ontology (GO) enrichment analysis

To identify candidate genes within the haploblocks of interest, a comprehensive functional annotation enrichment analysis was conducted using gene ontology (GO) information. To begin, the top 0.5% of haploblocks identified in the local GEBV analysis were selected based on the haploblocks with the highest variance for block effects (Aldiss et al. 2025). Physical positions of the markers defining these blocks were used to categorise genomic intervals. MorexV1 reference genome was the foundational assembly for the marker data and was used for the physical coordinates of each haploblock and the gene accessions within each interval were extracted. The list of identified genes underwent a GO enrichment analysis using the clusterProfiler package in R (Xu et al. 2024). This process involved building a custom gene annotation database from a GFF3 file for the MorexV1 (Hvulgare 462 r1) assembly and mapping gene identifiers to GO terms using an annotation file (Goodstein et al. 2012). A singular enrichment analysis was employed to determine if any GO terms were over-represented in the gene list from the haploblocks compared to the entire barley genome. The significance of enrichment was assessed using the Fisher’s Exact Test with a Benjamini-Hochberg false discovery rate correction (*p <* 0.05) to account for multiple testing. The identified genes were further validated and refined by cross-referencing against the updated MorexV3 assembly (Mascher et al. 2021). This was done using a combination of custom annotation files and the biomaRt package to query the Ensembl Plants database to ensure the most up to date gene information (Durinck et al. 2009; Dyer et al. 2025). Finally, the results were visualised using ggplot2 to create dot plots illustrating the most significantly enriched GO terms providing insight into the biological pathways associated with the traits of interest (Wickham 2016).

## 3 Results

### 3.1 Environments cluster by water stress profiles

The multi-environment trials (MET) assessed in this study encompassed eight environments (trial by year combination) across Western Australia and Queensland, which were categorized into low, moderate or high stress-intensity groups based on APSIM-simulated water stress indices (WSI). These environments represented a sample of a breeding program TPE ranging from low to high water stress conditions (figure S2).

Low-stress environments (GAT22, YRK22, and DND22) maintained high water availability throughout the growing season (mean WSI = 0.95 ± 0.06). The genotypic Best Linear Unbiased Predictions (GBLUPs) for these sites reflected similar yield levels across environments, with mean yields ranging from 4.39 t/ha at Dandaragan to 4.71 t/ha at Gatton (table 2). In contrast, moderate-stress environments (DND21 and GAT24) exhibited significant variability in water availability (SD = 0.17), particularly during the vegetative phase. This variability was reflected in the yield outcomes, with GAT24 producing the lowest mean yield of the MET at 3.68 t/ha and the highest coefficient of variation (6.61%).

**Table 2.** Summary of trial environments, stress classifications, and yield GBLUP statistics (t/ha)

| Site | Stress Group | $\mu_y$ | Min<sub>y</sub> | Max<sub>y</sub> | SE | CV (%) | Rep |
| --- | --- | --- | --- | --- | --- | --- | --- |
| CGN21 | High | 4.65 | 3.64 | 5.42 | 0.21 | 4.4 | 1.31 |
| CGN22 | High | 4.56 | 3.67 | 5.38 | 0.25 | 4.7 | 1.28 |
| DND21 | Moderate | 4.3 | 2.94 | 5.47 | 0.31 | 7.58 | 1.32 |
| DND22 | Moderate | 4.39 | 3.47 | 5.5 | 0.25 | 4.95 | 1.3 |
| GAT22 | Low | 4.71 | 4.23 | 5.09 | 0.12 | 2.08 | 1.62 |
| GAT24 | Moderate | 3.68 | 2.31 | 5.02 | 0.44 | 6.61 | 3.17 |
| YRK21 | High | 4.7 | 3.48 | 5.66 | 0.21 | 5.27 | 1.31 |
| YRK22 | Low | 4.45 | 3.65 | 5.15 | 0.22 | 4.26 | 1.3 |

$\mu_y$ = Site mean yield (t/ha); Min<sub>y</sub> = Minimum predicted yield; Max<sub>y</sub> = Maximum predicted yield; SE = Average standard error of prediction; CV = Coefficient of variation; Rep = mean level of replication.

High-stress environments, comprising CGN21, CGN22, and YRK21, were characterized by severe terminal drought stress (mean WSI = 0.61) and cooler mean temperatures (12.97 ◦C) relative to the other trial sites. In these environments, the WSI decreased significantly from 0.95 prior to flowering to approximately 0.30 by 200 ◦Cd post-flowering. Despite the rapid onset of terminal stress, these environments maintained competitive mean yields between 4.56 and 4.70 t/ha (table 2).

### 3.2 Genetic correlations reveal strong crossover *G* × *E* interaction

Analysis of grain yield across the eight environments revealed significant G×E interaction, with genetic correlations (*r*_*g*_) between pairs of environments ranging from strongly positive to moderately negative (figure 1A). For example, a strong positive correlation was observed between the Dandaragan environments across two seasons (DND21 and DND22; *r*_*g*_=0.89) indicating that genotype rankings were largely consistent and G×E was primarily non-crossover (scalar) in nature. In contrast, strong crossover (non-scalar) G×E was evident between other environments, such as GAT22 and the Corrigin environments (GAT22 vs. CGN21, *r*_*g*_ = −0.27; GAT22 vs. CGN22, *r*_*g*_ = −0.20). This negative correlation signifies that there is significant reranking of genotype performance in GAT22 compared to CGN environments, highlighting the complex G×E patterns present in this study.

**Fig. 1.**
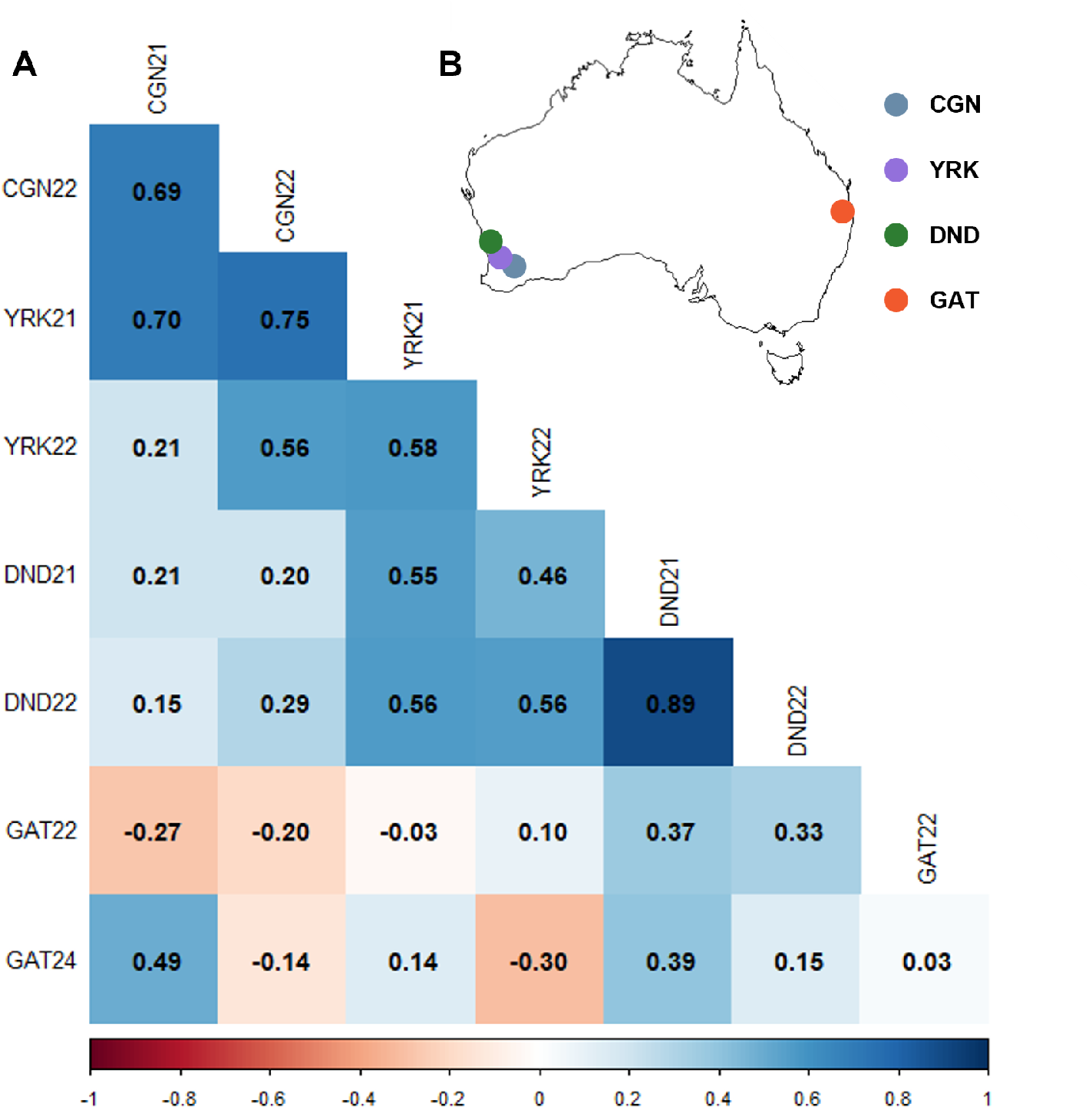
Overview of the multi-environment trial (MET) network and genetic relationships between environments. A) Correlogram of the additive genetic correlations between all pairs of environments. Genetic correlations were estimated from the additive genetic variance-covariance matrix of a factor analytic model of order three (FA3). The colour intensity and shade indicate the strength and direction of the correlation, with dark blue representing strong positive correlations and dark red indicating strong negative correlations, which signify crossover genotype-by-environment (G×E) interaction. B) Geographic locations of the eight field trials conducted in Western Australia (WA) and Queensland (QLD), Australia. Each point represents a unique environment, defined as a specific site-year combination. Trial site codes are: CGN (Corrigin, WA), YRK (York, WA), DND (Dandaragan, WA), and GAT (Gatton, QLD).

### 3.3 Genetic correlations reveal strong crossover *G* × *E* interaction

To model the *G* × *E* variance-covariance structure, an FA model of order three was fit to the additive genetic data partitioned through the genomic relationship matrix. The FA model successfully captured 80.47% of the total additive genetic variance across all eight environments. The model provided a robust fit for all environments, with the common factors explaining 100% of additive genetic variance for CGN21, DND21 and GAT24 (table 3). This was followed closely by CGN22, DND22 and YRK21 with 85.57, 85.12 and 81.43% respectively. However, only 26.51% of additive genetic variance was explained for GAT22, indicating this environment likely experienced unique conditions. Interestingly, while FA1 explained the majority of the genetic variation across most environments, for CGN21, CGN22 and GAT24 the main driver was FA2 and FA3 for GAT24 indicating magnitude differences in the strength of the *G* × *E* interactions.

**Table 3.**
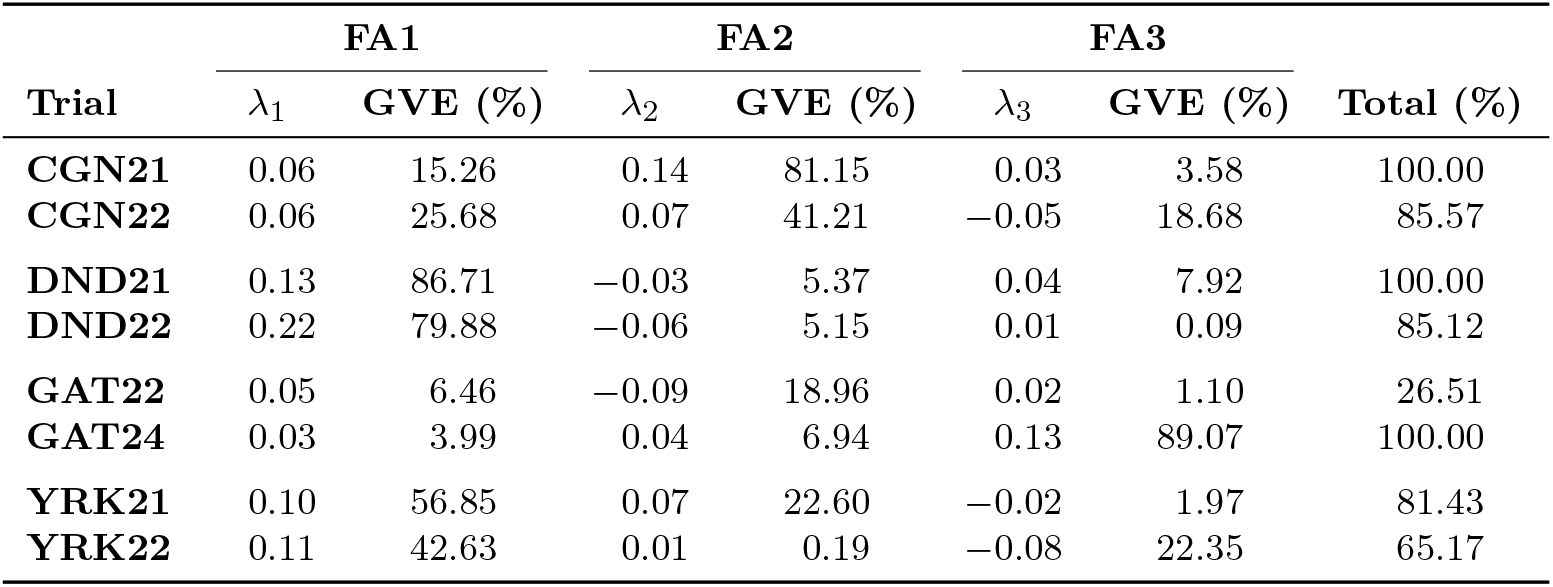
Factor analytic decomposition of additive genetic variance showing environmental loadings (*λ*) and genetic variance explained (GVE, %)

| Trial | FA1 |  | FA2 |  | FA3 |  | Total (%) |
| --- | --- | --- | --- | --- | --- | --- | --- |
| | $\lambda_1$ | GVE (%) | $\lambda_2$ | GVE (%) | $\lambda_3$ | GVE (%) | |
| CGN21 | 0.06 | 15.26 | 0.14 | 81.15 | 0.03 | 3.58 | 100.00 |
| CGN22 | 0.06 | 25.68 | 0.07 | 41.21 | -0.05 | 18.68 | 85.57 |
| DND21 | 0.13 | 86.71 | -0.03 | 5.37 | 0.04 | 7.92 | 100.00 |
| DND22 | 0.22 | 79.88 | -0.06 | 5.15 | 0.01 | 0.09 | 85.12 |
| GAT22 | 0.05 | 6.46 | -0.09 | 18.96 | 0.02 | 1.10 | 26.51 |
| GAT24 | 0.03 | 3.99 | 0.04 | 6.94 | 0.13 | 89.07 | 100.00 |
| YRK21 | 0.10 | 56.85 | 0.07 | 22.60 | -0.02 | 1.97 | 81.43 |
| YRK22 | 0.11 | 42.63 | 0.01 | 0.19 | -0.08 | 22.35 | 65.17 |

The rotated factor loadings were used, reflecting the commonalities in *G* × *E* across the MET (Smith and Cullis 2018). The first factor (*λ*_1_) had positive loadings across all environments, supporting its role as the primary source of additive genetic variance for yield mean performance (table 3). The magnitude of the loading reflects the strength of association with the trial and this latent factor, with DND22 having the strongest loading (*λ*_1_=0.22). Factors 2 (FA2 = *λ*_2_) and 3 (FA3 = *λ*_3_) captured the remaining G×E interactions. Factor 2 defined a strong contrast between CGN21 (*λ*_2_= 0.14) and GAT22 (*λ*_2_= -0.09), identifying the main axis of crossover interaction between these sites. Similarly, factor 3 primarily distinguished GAT24 (*λ*_3_ = 0.13) from YRK22 (*λ*_3_= -0.08), representing a different axis of G×E. Genotypes with high positive scores for Factor 2 would be expected to be adapted to CGN21 but poorly adapted to GAT22, while those with high positive scores for Factor 3 would be specifically adapted to GAT24.

### 3.4 Latent factor 2 is strongly correlated with water stress around flowering

To identify the environmental drivers of the G×E interaction, the FA loadings were correlated with the APSIM-simulated covariates. This analysis revealed a strong negative correlation between the FA2 loadings and the WSI during the thermal time window centred on flowering (*r* = −0.83; figure 2B). This establishes a clear biological basis for the main axis of G×E as FA2 quantitatively distinguishes environments based on their degree of water stress at this critical developmental stage. Environments with positive FA2 loadings (e.g. CGN21) experienced high water stress (low WSI; figure S2), while those with negative loadings (e.g. GAT22) had high water availability.

**Fig. 2.**
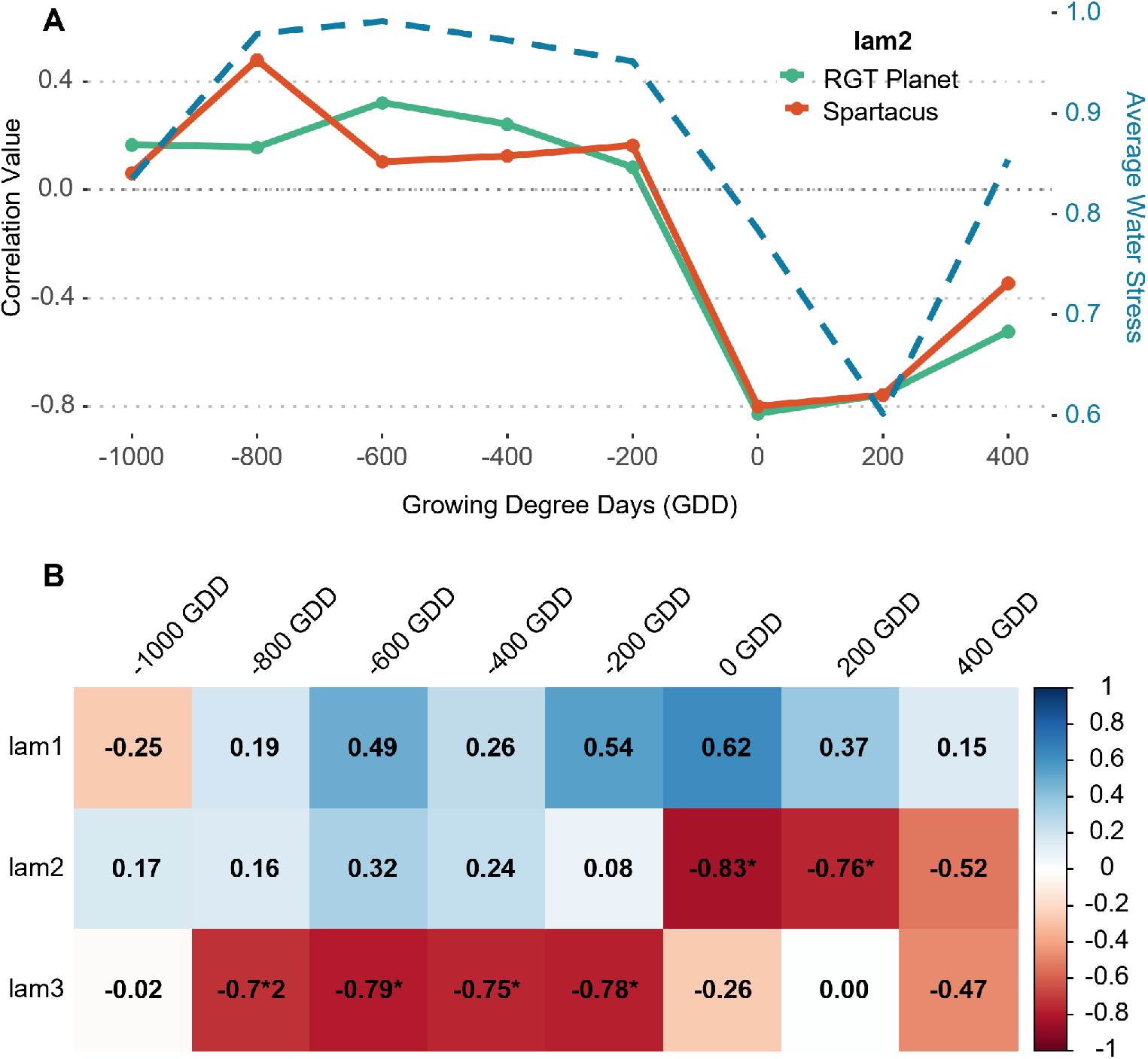
Environmental factor loadings correlate with water stress index. A) Correlation of lam2 environmental factor loading with WSI across GDD centered around flowering for RGT Planet (orange) and Spartacus (green). Average WSI across environments was fit for comparison. B) Pearson correlation matrix between the factor loadings (*λ*_1_, *λ*_2_, *λ*_3_) of the FA3 model and the APSIM simulated water stress index (WSI) at different thermal time points relative to flowering (GDD). Note that the smaller the WSI the more severe. WSI of 1.0 is considered high rainfall or water available. Red indicates a negative correlation and blue indicates positive. ∗ = *p <* 0.05

In addition, FA1 which represents overall yield performance, was moderately correlated with WSI around flowering (*r* = 0.62), indicating that higher-yielding environments generally experienced less water stress. FA3 showed only weak correlations at this key growth stage, though exhibited strong negative correlations with water stress index before flowering suggesting a potential relationship between WSI during development and the *G* × *E* patterns exhibited in this data set (figure 2B).

### 3.5 Haplotype Mapping Identifies Major Genomic Regions for Stability and Stress Response

Haplotype mapping was performed to identify genomic regions associated with yield potential represented by the latent *λ*1 and water stress response (WSR) corresponding to flowering by latent *λ*2. The local GEBVs for each haplotype were used to quantify effects of these two traits. Block variance was used to identify the blocks explaining the greatest variance for each trait and the top 3 blocks by variance were explored (figure 3).

**Fig. 3.**
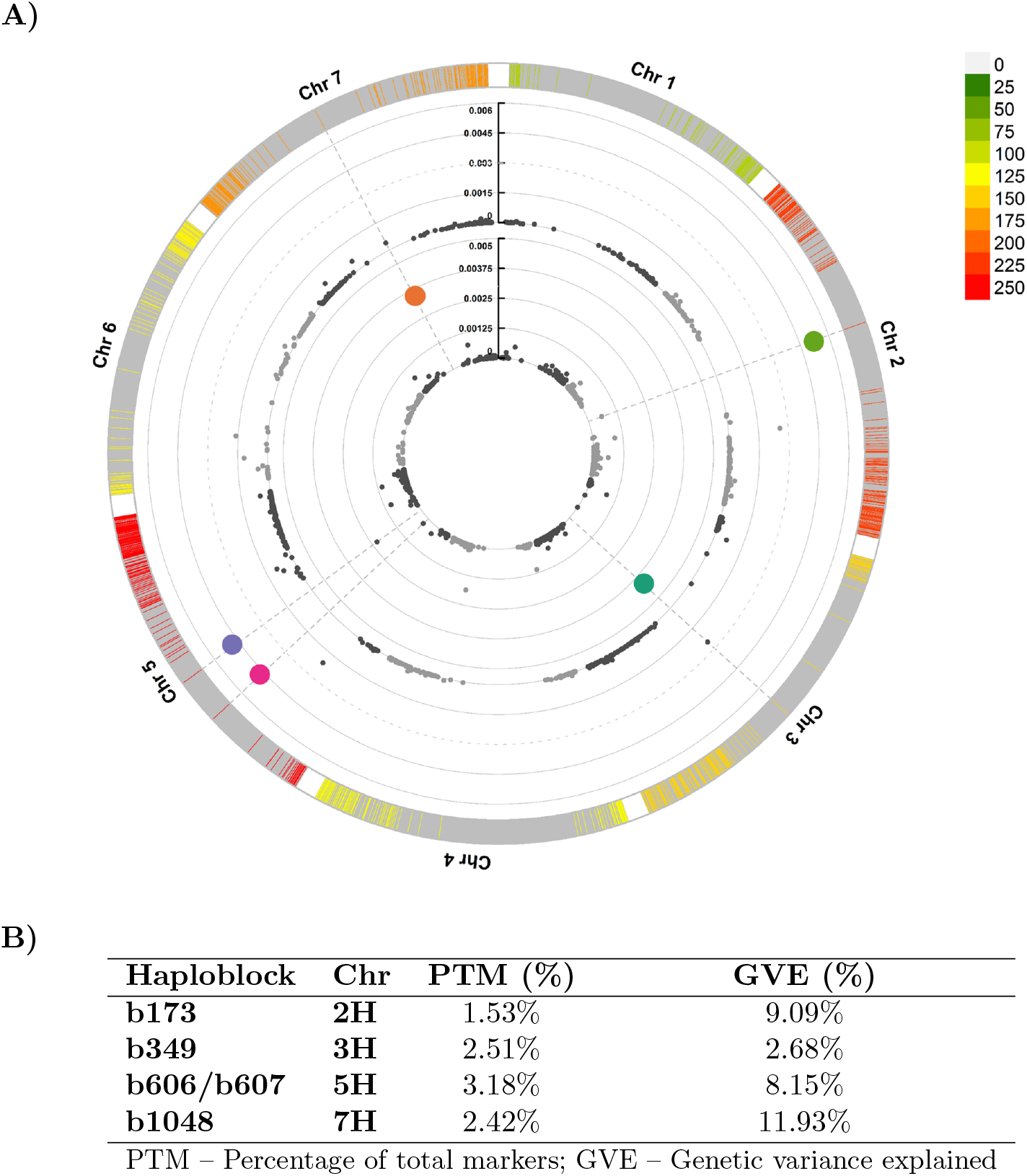
Haploblock-based mapping identifies major genomic regions associated with Yield Stability (FA1) and Water Stress Response (FA2). A) A circus plot highlighting the variance of haploblocks for 1,145 haploblocks across the chromosomes. The outer track displays marker density in 1 Mb windows. The inner scatter plot shows the variance of each haploblock’s effect, with higher values indicating greater contribution to the trait genetic variance. Dashed concentric circles represent variance thresholds. The top 0.5% of high-variance haploblocks are highlighted for Yield Stability (FA1, green), Water Stress Response (FA2, blue), or both traits (red). Key identified haploblocks include b349 (3H) and b1048 (7H) for FA1, and b173 (2H) and b607 (5H) for FA2. **B)** Summary of key haploblocks showing the percentage of total markers (PTM) and genetic variance explained (GVE).

For OP, a wide range of haplotype effects were observed with effect values ranging from -0.179 to 0.165 (Supplementary Data 1). The highest variance haploblock was b349 on chromosome 3H (at 355.68 cM). This block, containing 169 SNPs, was associated with the highest variance for this yield potential (0.004). The haplotype effects ranged from the most negative observed haplotype of -0.179 to 0.060. In contrast, b1048 on 7H (at 296.75 cM), exhibited the largest positive effect for OP, with values reaching 0.165. This block contained 163 SNP markers with a variance of 0.003.

For WSR, the haplotype block with the highest variance (0.005) was b173, located on chromosome 2H at a genetic position of 328.44 cM. This block, which comprised 103 SNPs, showed a range of haplotype effects from -0.055 to 0.152 across genotypes, accounting for the largest positive effect on water stress response. Additional major effects were found on chromosome 5H, b000606 (at 161.7 cM), contained 142 SNPs and with a variance of 0.0046 consisted of haplotype effects ranging from -0.003 to -0.185, indicating a negatively skewed association with both positive and negative effects on the WSR trait. Adjacent to b606 is b607 the only block identified across both analyses; the second highest variance for OP and third highest for WSR with a variance of 0.002 and 0.005 respectively. The block contains containing 72 SNPs exhibited the largest negative effect of -0.215 for WSR and second largest positive effect for OP of 0.144.

To explore the contribution of each of these blocks the percentage of genetic variance they accounted for was explored and compared to the percentage of markers the block contained. Despite being the highest variance block for OP b349 accounted for an amount of additive genetic variance almost proportional to the number of markers it contained. In contrast b1048 accounted for 11.93% of additive genetic variance despite having less markers than b349. As blocks b606 and b607 were adjacent, it was determined there would be too much colinearity to calculate their variance contributions independently and thus were combined in this analysis accounting for 8.15% of the additive genetic variance for yield.

### 3.6 Distribution of Favourable Haplotypes in Elite Cultivars

To translate the haploblock mapping into actionable breeding knowledge, the distribution of Haplotypes of Interest (HOIs) at key high-variance loci was explored. This was done within four elite commercial cultivars, Banks, RGT Planet, Maximus CL, and Scope CL selected for their distinct combinations of favourable alleles (table S5). This analysis aimed to determine the prevalence of favourable alleles in modern germplasm and to understand how these alleles combine in different genetic backgrounds. For each haploblock, effects were scaled to a mean of zero, ensuring that positive values consistently represent a desirable genetic effect, even for traits where all haplotypes might have a net-negative impact.

For OP, the distribution of haplotype effects differed markedly between the two major loci (figure 4A). Haploblock b349 exhibited a bimodal distribution, with a primary mode centered on the population mean and a smaller, favourable mode approximately two standard deviations above it. Haplotypes within this positive mode were classified as HOIs and were present in both Banks and RGT Planet. Haploblock b607 exhibited a similar pattern with two modes the larger of which having a positive scaled effect once again both Banks and RGT planet had HOI at this block. In contrast, Haploblock b1048 showed a strongly skewed distribution where most haplotypes had the same scaled positive effect, indicating that favourable alleles at this locus are common in the elite population. Consequently, only Scope CL lacked an HOI at this block. Reflecting the sum of these and other loci, the overall genetic merit (BLUP) for yield potential was highest in Maximus CL and lowest in Scope CL among the four cultivars (table S5).

**Fig. 4.**
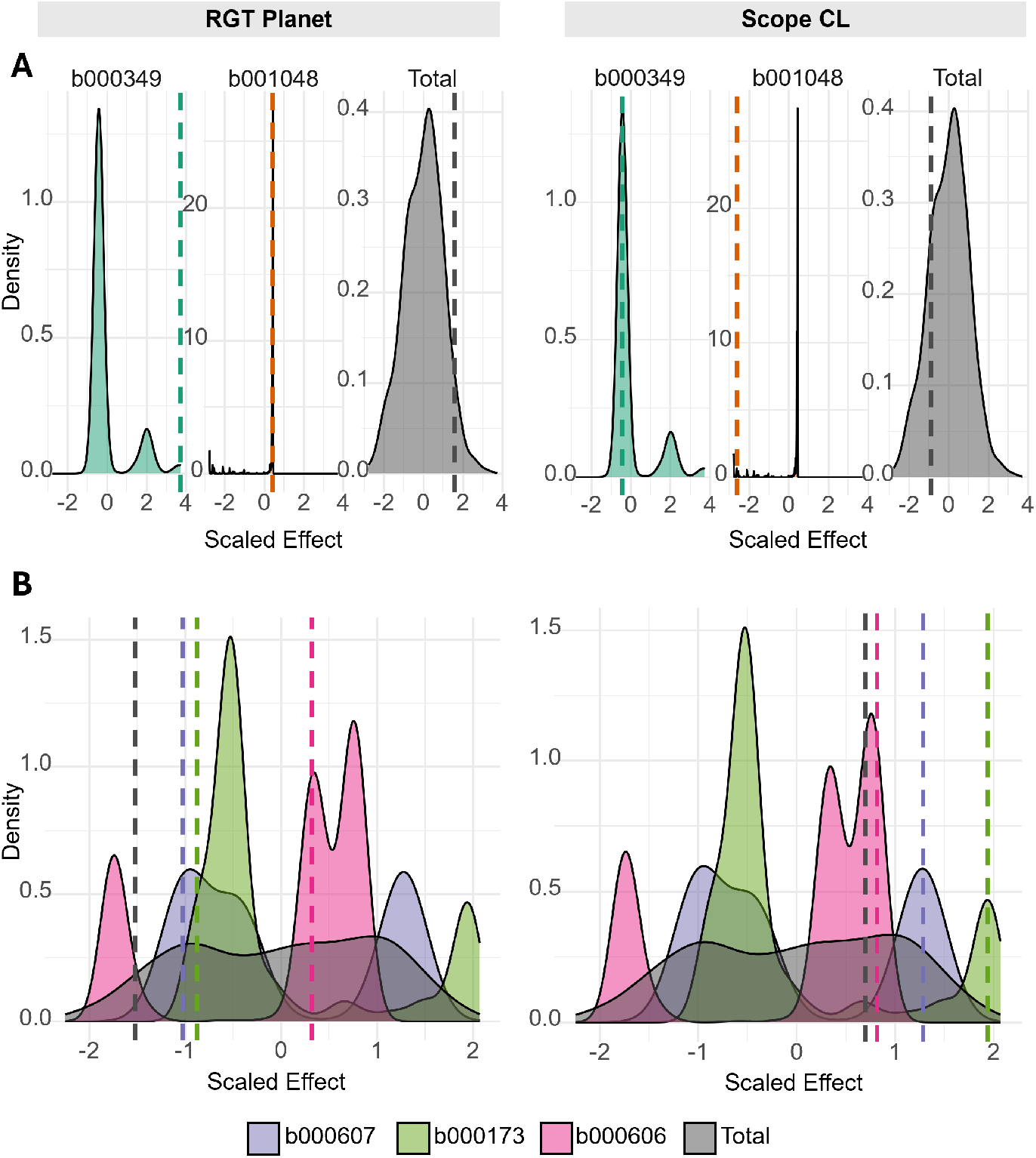
Distribution of scaled haplotype effects for major haploblocks associated with Yield Stability and Water Stress Response in elite barley cultivars RGT Planet and Scope CL. Density plots illustrate the distribution of scaled local GEBV effects for all haplo-types present in the population for key haploblocks. The vertical black line indicates the specific scaled effect of the haplotype carried by the named cultivar. Plots for haploblocks b349 and b1048, and the total GEBV for Yield Stability (FA1). A) Plots for haploblocks b349 and b1048, and the total GEBV for Yield Stability (FA1). B) Plots for haploblocks b607, b173, and b606, and the total GEBV for Water Stress Response (FA2). The x-axis represents the scaled effect, where the population mean is zero. These plots reveal the genetic architecture of adaptation in key cultivars and the distribution of favourable and unfavourable alleles within the broader breeding population.

The haploblocks associated with WSR also showed distinct patterns of allelic distribution (figure 4B). Haploblock b173 displayed a bimodal distribution, but the favourable mode was rarer and had a larger effect as both Banks and Scope CL carried a highly desirable haplotype with an effect more than two standard deviations above the population mean. For Haploblock b607 the effects were distributed similarly to the OP analysis. Haploblock b606 had a multi-model distribution with three distinct peaks one of which contained negative haplotypes at two standard deviations below the mean. Interestingly the adjacent Haploblocks b606 and b607 on chromosome 5H appeared to be inherited together with Maximus CL and Scope CL carried HOIs for both loci. Overall, Maximus CL had the highest genetic merit for favourable WSR, while RGT Planet had the lowest.

### 3.7 Gene Ontology Enrichment Implicates Key Stress and Development Pathways

To infer the biological function of the high-variance genomic regions, a Gene Ontology (GO) enrichment analysis was performed on the genes residing within the highest variance haploblocks for each trait (table 4). For the OP trait, two primary regions were analysed. Haploblock b349 on chromosome 3H was significantly enriched for GO terms related to proton and ATP transport and homeostasis (figure S3). This region contained 226 genes, including putative candidates such as *V-type ATPase* subunits and the ATP-dependent metalloprotease *FTSH11*. The second region, haploblock b1048 on chromosome 7H, contained 219 genes and was enriched for GO terms associated with root and canopy development, with all significant hits encoding *Protein BPS1*.

**Table 4.**
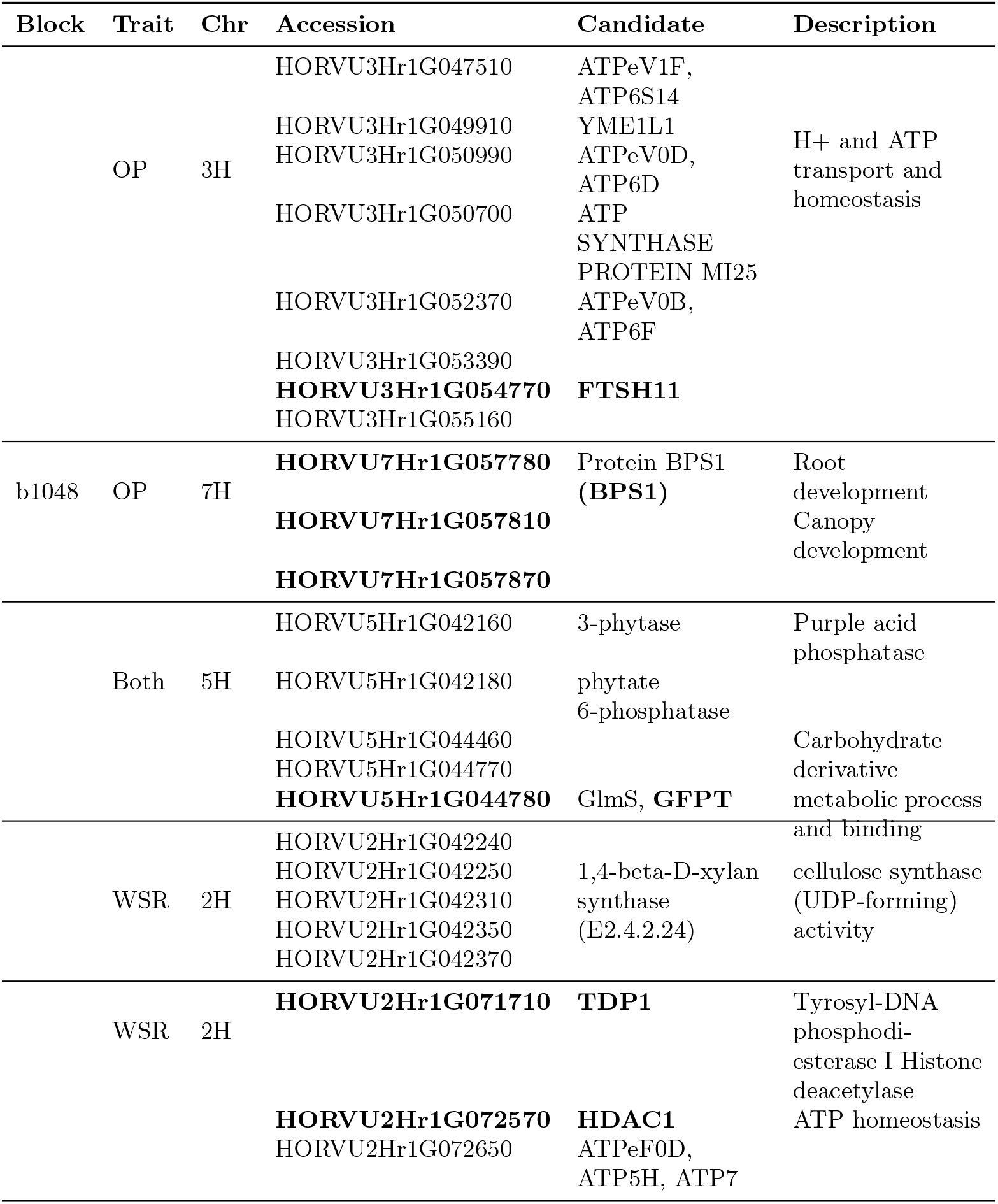
Haploblock summary and candidate genes for overall performance (OP) and water stress response (WSR) traits.

For the water stress response trait, two significant haploblocks were identified on chromosome 2H. Haploblock b173 showed enrichment for terms related to cellulose synthase activity, with a *1,4-beta-D-xylan synthase* gene as the primary candidate (figure S3). The adjacent Haploblock b177 was enriched for GO terms associated with ATP homeostasis and DNA damage response, containing candidate genes such as *tyrosyl-DNA phosphodiesterase I* (*TDP1*) and *HDAC1* (table 4). Finally, haploblock b607 on chromosome 5H, which was associated with both the OP and WSR traits, was enriched for genes involved in phosphate and carbohydrate metabolic processes, including *3-phytase* and *phytate 6-phosphatase* (table 4).

## 4 Discussion

This study presents a framework that bridges the predictive power of statistical *G* × *E* models with the explanatory power of functional genomics. The conceptual novelty is the treatment of genotypic scores from latent FA regression as novel and heritable quantitative traits. This innovation transforms qualitative statistical outputs into biologically meaningful metrics such as yield potential and yield stability under water stress. By establishing this essential link, we enable the use of powerful tools like haplotype-based mapping to dissect the genetic architecture of adaptation itself. This mapping technique is expanded with application of local GEBV calculations to further classify the main positive and negative drivers of yield with exploration into the desirable HOI and their presence within commercial cultivars. This approach moves beyond simply predicting which genotypes will perform best and begins to explain why, providing a clear path to identifying the underlying genetics and candidate genes driving environmental resilience. Similarly, with the environmental covariate it is possible to predict into new environments, beyond the MET into the TPE.

The targeted approach employed in this study involved correlating the environmental loadings derived from each latent factor of the FA-LMM with a suite of environmental covariates simulated by APSIM across different developmental stages. The resulting strong negative correlation (*r* = −0.84) between the FA2 environmental loadings and the WSI during the critical flowering period and early grain filling provides the central biological insight of the analysis (figure 2). This finding establishes a clear, quantitative link as the primary axis of crossover G×E observed in this elite barley population is driven by differential genotypic responses to water availability during flowering. Environments with positive loadings on FA2 were those that experienced high water stress (low WSI) at flowering, while environments with negative loadings were characterized by low water stress (high WSI).

Where current research focuses on the paradigm of capturing as much *G* × *E* variance as possible to remove from the model, this approach captures the *G* × *E* that is useful, interpretable and changeable within the population. Tolhurst et al. (2019) built on the principals of FA models by incorporating environmental covariates with latent factors to capture and partition scale and cross-over *G* × *E* (Tolhurst et al. 2019). However, fitting these environmental covariates directly poses a challenge for both interpretation and extrapolation into future environments (Piepho and Williams 2024). Instead of fitting these values directly, through crop growth models it is possible to identify and validate statistical patterns using underlying biology (Powell et al. 2022).

The genetic architecture of *G* × *E* response traits offers a glimpse into the history of Australian barley breeding. High variance haplotypes for the candidate haploblocks were mostly specific to either OP or WSR with one overlap at b607 (figure 3). The amount of additive genetic variance attributed to these blocks was greater than their percentage of the total markers further indicating their role in yield. The fact that the HOIs are already well-distributed within this elite material suggests that breeders have been inadvertently selecting for these adaptive mechanisms through routine, phenotype-based evaluation in Australia’s variable production environments. These patterns likely represent distinct haplotype families, perhaps originating from different founder germplasm or reflecting divergent selection pressures for competing objectives (e.g. adaptation to drought vs high-yielding potential). This is evidenced by the HOIs for b607 being opposites for OP and WSR, for example haplotypes Banks and RGT Planet have positive scaled effects for OP and negative for WSR (figure 4A, B). This suggests that major historical breeding decisions have left a detectable signature on the genetic architecture of environmental adaptation. This approach of defining HOIs based on these population distributions, rather than a simple effect threshold, provides breeders with a broader set of validated, superior alleles, facilitating selection while helping to manage genetic diversity (Brunner et al. 2024; Vo Van-Zivkovic et al. 2025).

While statistically significant, the identification of candidate genes is based solely on functional annotation, which does not establish causality. Where the haplotype-based mapping approach successfully narrowed down large genomic regions to high-variance haploblocks, the most immediate next step is the functional validation of the identified candidate genes contributing to the block effects. Similarly, expanding this approach in a bigger population to capture the complete TPE will provide more refined selection of gene targets. Even so, the distillation of haploblocks to a handful of individual genes identified several candidates with functional annotations that seemingly coincided with the assessed traits. For example, *Filamentous temperature sensitive H (FTsH11)* and *Bypass1 (BPS1)* were identified as notable genetic drivers within the highest variance haploblocks contributing to OP. Both genes have been functionally dissected in model plant species where FTsH11 has shown to be crucial in modulating photosynthetic efficiencies under temperature stress while BPS1 has been shown to play a role in long distance signalling between the root and shoot (Chen et al. 2006; Van Norman et al. 2004). Similarly, TDP1 was found to coincide with one of the highest variance haploblocks for FA2 (water stress response), which has been reported to play a key role in programmed cell death, a process known to be involved in drought stress (Lee et al. 2010; Shamloo-Dashtpagerdi et al. 2020). Modern gene-editing technologies, particularly CRISPR/Cas9, provide the ideal tools for further dissecting and beneficially mutating the identified genes. By creating targeted knockout mutants or allele-swap lines for genes such as FTSH11, BPS1, and TDP1 in an elite barley background, their precise contribution to yield potential and water stress response can be unequivocally tested under controlled-environment and field conditions. Such experiments would provide the ultimate proof of function and could accelerate the direct engineering of superior alleles into breeding lines.

## 5 Conclusion

The flexibility of this framework was demonstrated through the integration of latent environmental covariates. While the analysis was limited to eight trials, the results indicated that specific genetic drivers for a TPE can be identified, suggesting the potential utility of this approach for larger datasets. Although the analysis was specific to the MET assessed in this study, the methodology provides a characterisation system applicable to other breeding programs. Any breeding program can use its MET data to identify the unique environmental pressures driving G×E in its specific target environments, map the relevant adaptive loci, and develop tailored selection strategies. This framework offers a potential refinement of selection strategies, moving from the use of a single, complex genomic estimated breeding value for yield towards the identification of underlying functional components. The methodology presented here dissects the ‘black box’ of yield into its functional constituents. The dissection of yield into functional constituents allows for the future identification of markers associated with specific physiological mechanisms such as energy homeostasis or plant signalling which contribute to resilience within a TPE. This transition towards mechanism-based selection enables the rational design of cultivars tailored for future climates, promising more rapid and reliable genetic gain.

## Acknowledgments

The authors are grateful for the technical and breeding staff at InterGrain Pty. Ltd for providing the germplasm and genotypic data used in this study and thanks to Sarah van der Meer and Samir Alahmad at the University of Queensland for running the Gatton trials.

## Declarations

### Funding

This research was funded by the Australian Research Council Industry Linkage Project ‘Digging Deeper to Improve yield potential’ (LP200200927) in collaboration with InterGrain Pty. Ltd. LH was supported through an ARC Future Fellowship (FT220100350). YL, ZA, SMB were supported as an Affiliated PhD and Postdocs of the Australian Research Council Training Centre in Predictive Breeding for Agricultural Futures (IC230100016). BH, SMB and ZA received scholarships from the Australian Government Research Training Program. ZA received scholarship from InterGrain. SMB was supported by Deutsche Forschungsgemeinschaft (DFG) and University of Queensland IRTG 2843 – Accelerating Crop Genetic Gain.

### Conflict of interest statement

The authors have no relevant financial or non-financial interests to disclose.

### Data availability statement

The data that support the findings of this study are available from InterGrain Pty Ltd, but restrictions apply to the availability of these data, which were used under licence for the current study, and so are not publicly available. Data are however available from the authors upon reasonable request and with permission of InterGrain Pty Ltd.

### Code availability

Code is available upon request to authors.

### Author contributions

ZA and YL conceived the idea and study. ZA performed MET analysis and latent regression. SMB performed haplotype analysis. ZA and YL identified candidate genes and performed block level GO enrichment. SB, DM and DG provided the data. BH and KC performed envirotyping and APSIM modelling. ZA, SMB, and YL wrote the manuscript. ZA and SMB designed the figures. LH and YL managed the project. All authors read, edited, and approved the final manuscript.

## Supplementary Materials

**Fig. S1.**
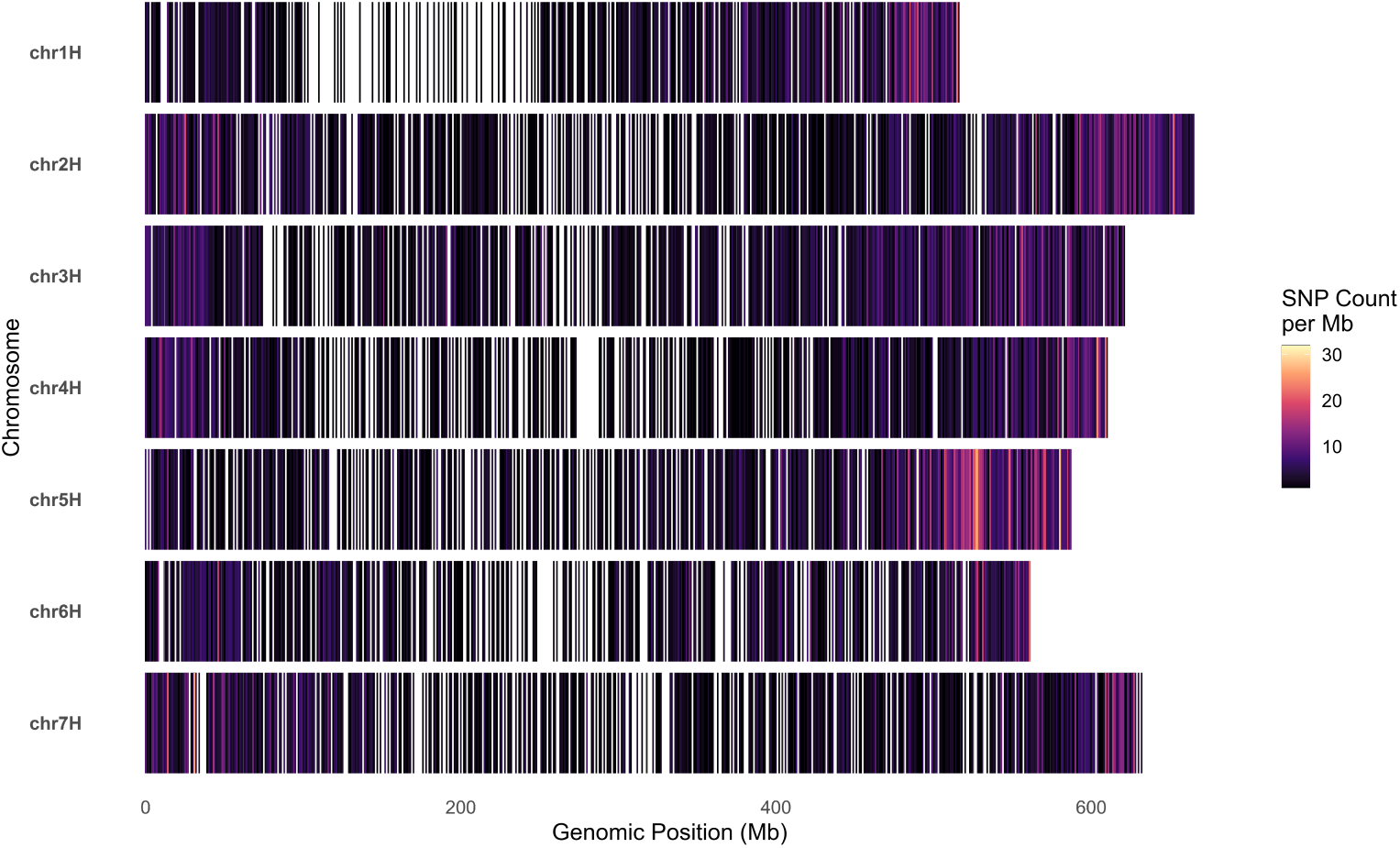
SNP marker density heatmap. SNP marker density of the final 6,733 high quality SNP markers presented on the MorexV3 Genome. Density represented by colour intensity in 1 Mb windows.

**Fig. S2** Characterisation of environmental conditions across trial eight sites

Environmental covariates were simulated using the Agricultural Production Systems sIMulator (APSIM) for each of the eight environments. Panels display the daily (A) Water Stress Index (WSI), (B) mean temperature (°C), (C) maximum temperature (°C), and (D) minimum temperature (°C). Data are plotted against thermal time in growing degree days (°Cd) relative to the simulated flowering date (0 °Cd). Each coloured line represents a unique trial-year combination, as indicated in the legend. A WSI value approaching 1.0 indicates minimal water stress, whereas a value approaching 0 indicates severe water stress.

**Fig. S3.**
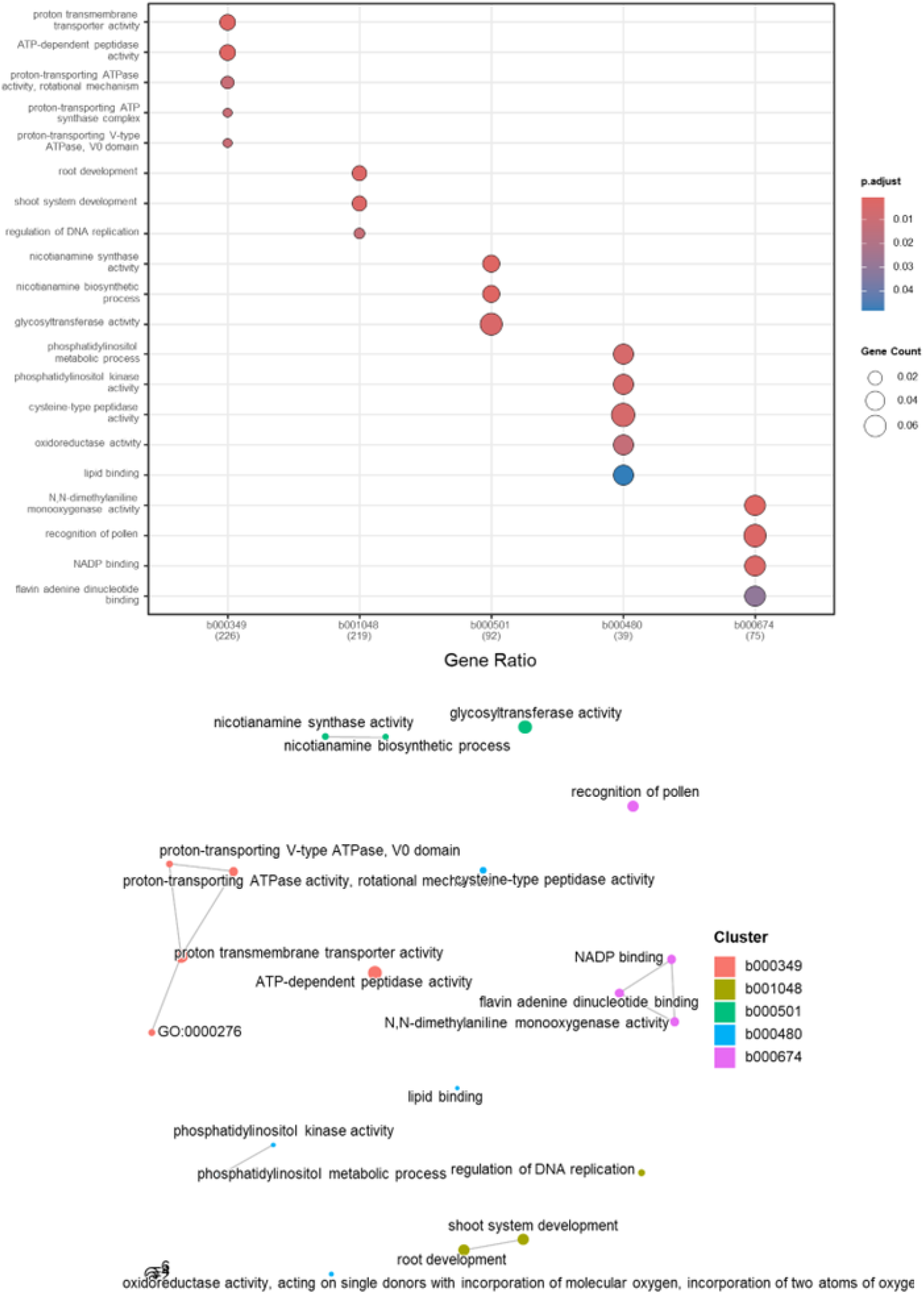
Gene Ontology (GO) enrichment analysis reveals biological functions of candidate genes within high-variance haploblocks for FA1. (A) Functional relationship map of enriched GO terms for haploblocks associated with Yield Stability (FA1). Nodes represent significantly enriched GO terms, with colors indicating the haploblock cluster. (B) Dot plot summarising the GO enrichment results for FA1. The x-axis represents the gene ratio (proportion of genes in the haploblock associated with a term), dot size represents the number of genes (Gene Count), and dot color indicates the adjusted p-value.

**Fig. S4** Gene Ontology (GO) enrichment analysis reveals biological functions of candidate genes within high-variance haploblocks for FA2

(A) Functional relationship map for haploblocks associated with Water Stress Response (FA2). (B) Dot plot summarising the GO enrichment results for FA2. Key enriched pathways for FA1 included ATP homeostasis and root development, while pathways for FA2 included cellulose synthesis and defense response.

**Table S1.** Summary of trial sites, including unique genotypes and replication percentages.

| Site | Total<br>Unique | Genotyped | Ungenotyped | Pct. Rep.<br>(Indiv.) | Pct. Rep.<br>(Across) |
| --- | --- | --- | --- | --- | --- |
| CGN21 | 734 | 458 | 276 | 26.4 | 100.0 |
| CGN22 | 737 | 643 | 94 | 27.1 | 100.0 |
| DND21 | 734 | 458 | 276 | 26.4 | 100.0 |
| DND22 | 737 | 643 | 94 | 27.1 | 100.0 |
| GAT22 | 398 | 338 | 60 | 43.0 | 99.5 |
| GAT24 | 267 | 216 | 51 | 55.4 | 98.1 |
| YRK21 | 734 | 458 | 276 | 26.4 | 100.0 |
| YRK22 | 737 | 643 | 94 | 27.1 | 100.0 |
| YRK24 | 267 | 216 | 51 | 55.4 | 98.1 |

**Note:** Pct. Rep. (Indiv.) = percentage of genotypes replicated within the individual site; Pct. Rep. (Across) = percentage of genotyped lines present at the site that are replicated elsewhere in the trial network.

**Table S2.** Genotype concurrence matrix (%) across trial sites.

|  | CGN21 | CGN22 | DND21 | DND22 | GAT22 | GAT24 | YRK21 | YRK22 |
| --- | --- | --- | --- | --- | --- | --- | --- | --- |
| **CGN21** | 100.0 | 67.0 | 100.0 | 67.0 | 33.4 | 22.1 | 100.0 | 67.0 |
| **CGN22** | 66.8 | 100.0 | 66.8 | 100.0 | 53.7 | 27.0 | 66.8 | 100.0 |
| **DND21** | 100.0 | 67.0 | 100.0 | 67.0 | 33.4 | 22.1 | 100.0 | 67.0 |
| **DND22** | 66.8 | 100.0 | 66.8 | 100.0 | 53.7 | 27.0 | 66.8 | 100.0 |
| **GAT22** | 61.6 | 99.5 | 61.6 | 99.5 | 100.0 | 25.9 | 61.6 | 99.5 |
| **GAT24** | 60.7 | 61.5 | 60.7 | 61.5 | 38.6 | 100.0 | 60.7 | 61.5 |
| **YRK21** | 100.0 | 67.0 | 100.0 | 67.0 | 33.4 | 22.1 | 100.0 | 67.0 |
| **YRK22** | 66.8 | 100.0 | 66.8 | 100.0 | 53.7 | 27.0 | 66.8 | 100.0 |

**Note:** Values represent the percentage of shared genotypes between trial sites (mean off-diagonal concurrence = 67.7%). Diagonals represent 100% self-concurrence.

**Table S3.** FA model statistics.

| Model | loglik | seq.lrt | df | pval | AIC | BIC | perc.var |
| --- | --- | --- | --- | --- | --- | --- | --- |
| *Single Site* |  |  |  |  |  |  |  |
| diag | 1386.78 | NA | NA | NA | -2685.56 | -2379.53 | NA |
| *G × E* |  |  |  |  |  |  |  |
| fa1 | 1860.09 | 946.62 | 7 | 0.00 | -3618.18 | -3263.47 | 38.46 |
| fa2 | 1897.58 | 74.98 | 7 | 0.00 | -3679.16 | -3275.76 | 52.24 |
| fa3 | 1904.45 | 13.73 | 6 | 0.03 | -3680.90 | -3235.76 | 58.26 |
| fa4 | 1908.60 | 8.31 | 4 | 0.08 | -3681.21 | -3208.25 | 77.97 |

**Note:** loglik = residual log-likelihood; seq.lrt = sequential likelihood ratio statistic (REMLRS); df = degrees of freedom; pval = p-value; AIC = Akaike Information Criterion; BIC = Bayesian Information Criterion; perc.var = average total genetic variance explained across sites. No GRM was included to determine best fit.

**Table S4.**
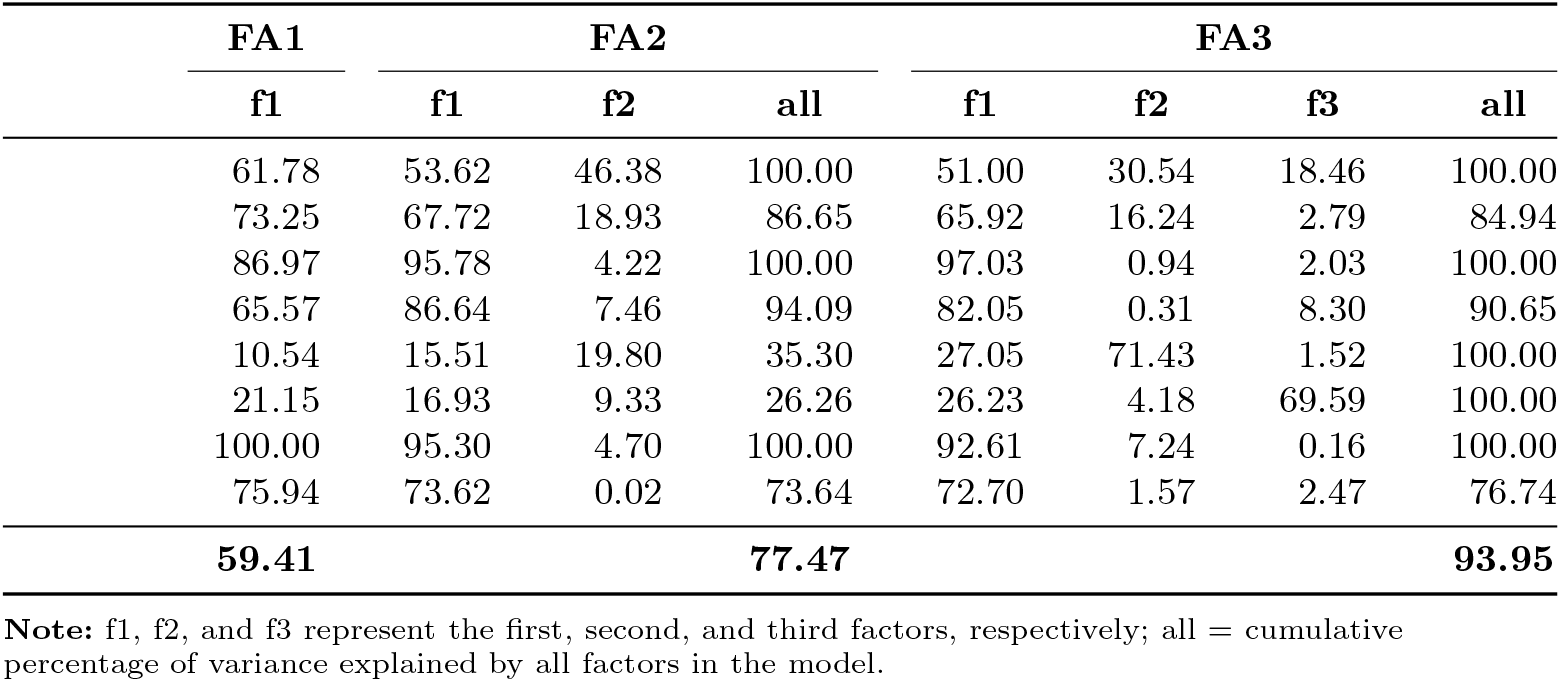
Percentage of genetic variance explained by individual factors and total for FA1, FA2, and FA3 models across sites.

| Site | FA1 | FA2 |  |  | FA3 |  |  |  |
| --- | --- | --- | --- | --- | --- | --- | --- | --- |
|  | f1 | f1 | f2 | all | f1 | f2 | f3 | all |
| CGN21 | 61.78 | 53.62 | 46.38 | 100.00 | 51.00 | 30.54 | 18.46 | 100.00 |
| CGN22 | 73.25 | 67.72 | 18.93 | 86.65 | 65.92 | 16.24 | 2.79 | 84.94 |
| DND21 | 86.97 | 95.78 | 4.22 | 100.00 | 97.03 | 0.94 | 2.03 | 100.00 |
| DND22 | 65.57 | 86.64 | 7.46 | 94.09 | 82.05 | 0.31 | 8.30 | 90.65 |
| GAT22 | 10.54 | 15.51 | 19.80 | 35.30 | 27.05 | 71.43 | 1.52 | 100.00 |
| GAT24 | 21.15 | 16.93 | 9.33 | 26.26 | 26.23 | 4.18 | 69.59 | 100.00 |
| YRK21 | 100.00 | 95.30 | 4.70 | 100.00 | 92.61 | 7.24 | 0.16 | 100.00 |
| YRK22 | 75.94 | 73.62 | 0.02 | 73.64 | 72.70 | 1.57 | 2.47 | 76.74 |
| **Total** | **59.41** |  |  | **77.47** |  |  |  | **93.95** |

**Note:** f1, f2, and f3 represent the first, second, and third factors, respectively; all = cumulative percentage of variance explained by all factors in the model.

**Table S5.** Presence or Absence of Haplotype of Interest in 12 barley cultivars.

| *HOI* | **FA1** |  | **FA2** |  |  |
| --- | --- | --- | --- | --- | --- |
|  | **b349** | **b1048** | **b607** | **b173** | **b606** |
| **Banks** | Yes | Yes | No | Yes | No |
| **Barque** | No | Yes | No | Yes | No |
| **Combat** | No | Yes | Yes | Yes | Yes |
| **Compass** | No | Yes | Yes | No | Yes |
| **Fairing** | No | Yes | Yes | No | Yes |
| **Laureate** | Yes | Yes | No | No | No |
| **Maximus CL** | No | Yes | Yes | No | Yes |
| **Neo CL** | Yes | Yes | No | No | No |
| **RGT** | Yes | Yes | No | No | No |
| **Rosalind** | No | No | No | No | No |
| **Scope CL** | No | No | Yes | Yes | Yes |
| **Spartacus CL** | No | Yes | Yes | No | Yes |

## Notes

### Competing Interest Statement

The authors have declared no competing interest.

## References

Aldiss Z, Lam Y, Baraibar S, et al (2025) Haplotype-based insights into seminal root angle in barley. The Plant Genome 18(3):e70088

Amin A, Christopher J, Cooper M, et al (2025) Envirotyping facilitates understanding of genotype× environment interactions and highlights the potential of stay-green traits in wheat. Field Crops Research 331:109940

Bančič J, Ovenden B, Gorjanc G, et al (2023) Genomic selection for genotype performance and stability using information on multiple traits and multiple environments. Theoretical and Applied Genetics 136(5):104

Brunner SM, Dinglasan E, Baraibar S, et al (2024) Characterizing stay-green in barley across diverse environments: unveiling novel haplotypes. Theoretical and Applied Genetics 137(6):120

Butler D, Cullis B, Gilmour A, et al (2017) ASReml-R reference manual version 4. VSN International Ltd, Hemel Hempstead, HP1 1ES, UK

Chen J, Burke JJ, Velten J, et al (2006) FtsH11 protease plays a critical role in Arabidopsis thermotolerance. The Plant Journal 48(1):73–84

Chenu K, Deihimfard R, Chapman SC (2013) Large-scale characterization of drought pattern: a continent-wide modelling approach applied to the Australian wheat-belt–spatial and temporal trends. New Phytologist 198(3):801–820

Durinck S, Spellman PT, Birney E, et al (2009) Mapping identifiers for the integration of genomic datasets with the R/Bioconductor package biomaRt. Nature protocols 4(8):1184–1191

Dyer SC, Austine-Orimoloye O, Azov AG, et al (2025) Ensembl 2025. Nucleic acids research 53(D1):D948–D957

Endelman JB (2011) Ridge regression and other kernels for genomic selection with R package rrBLUP. The plant genome 4(3)

Goodstein DM, Shu S, Howson R, et al (2012) Phytozome: a comparative platform for green plant genomics. Nucleic acids research 40(D1):D1178–D1186

Hill WG, Mackay TF (2004) DS Falconer and Introduction to quantitative genetics. Genetics 167(4):1529–1536

Holzworth D, Huth NI, Fainges J, et al (2018) APSIM Next Generation: Overcoming challenges in modernising a farming systems model. Environmental Modelling & Software 103:43–51

Holzworth DP, Huth NI, deVoil PG, et al (2014) APSIM–evolution towards a new generation of agricultural systems simulation. Environmental Modelling & Software 62:327–350

Jeffrey SJ, Carter JO, Moodie KB, et al (2001) Using spatial interpolation to construct a comprehensive archive of Australian climate data. Environmental Modelling & Software 16(4):309–330

Kelly AM, Smith AB, Eccleston JA, et al (2007) The accuracy of varietal selection using factor analytic models for multi-environment plant breeding trials. Crop Science 47(3):1063–1070

Lee SY, Kim H, Hwang HJ, et al (2010) Identification of tyrosyl-DNA phosphodiesterase as a novel DNA damage repair enzyme in Arabidopsis. Plant Physiology 154(3):1460–1469

Li X, Guo T, Wang J, et al (2021) An integrated framework reinstating the environmental dimension for GWAS and genomic selection in crops. Molecular Plant 14(6):874–887. 10.1016/j.molp.2021.03.010, URL https://www.sciencedirect.com/science/article/pii/S167420522100085X

Marschner IC, Schou IM (2019) Underestimation of treatment effects in sequentially monitored clinical trials that did not stop early for benefit. Statistical methods in medical research 28(10-11):3027–3041

Mascher M, Wicker T, Jenkins J, et al (2021) Long-read sequence assembly: a technical evaluation in barley. The Plant Cell 33(6):1888–1906

Oliveira ICM, Guilhen JHS, Ribeiro PCdO, et al (2020) Genotype-by-environment interaction and yield stability analysis of biomass sorghum hybrids using factor analytic models and environmental covariates. Field Crops Research 257:107929. 10.1016/j.fcr.2020.107929, URL https://www.sciencedirect.com/science/article/pii/S0378429020312132

Piepho H, Williams E (2024) Factor-analytic variance–covariance structures for prediction into a target population of environments. Biometrical Journal 66(6):e202400008

Powell OM, Barbier F, Voss-Fels KP, et al (2022) Investigations into the emergent properties of gene-to-phenotype networks across cycles of selection: a case study of shoot branching in plants. in silico Plants 4(1):diac006

Shaffer W, Papin V, Yadav S, et al (2025) Local genomic estimates provide a powerful framework for haplotype discovery. bioRxiv pp 2025–08

Shamloo-Dashtpagerdi R, Lindlöf A, Aliakbari M, et al (2020) Plausible association between drought stress tolerance of barley (Hordeum vulgare L.) and programmed cell death via MC1 and TSN1 genes. Physiologia plantarum 170(1):46–59

Smith A, Cullis B, Thompson R (2001) Analyzing variety by environment data using multiplicative mixed models and adjustments for spatial field trend. Biometrics 57(4):1138–1147

Smith AB, Cullis BR (2018) Plant breeding selection tools built on factor analytic mixed models for multi-environment trial data. Euphytica 214(8):143

Smith AB, Ganesalingam A, Kuchel H, et al (2015) Factor analytic mixed models for the provision of grower information from national crop variety testing programs. Theoretical and applied genetics 128(1):55–72

Tolhurst DJ, Mathews KL, Smith AB, et al (2019) Genomic selection in multi-environment plant breeding trials using a factor analytic linear mixed model. Journal of Animal Breeding and Genetics 136(4):279–300

Van Norman JM, Frederick RL, Sieburth LE (2004) BYPASS1 negatively regulates a root-derived signal that controls plant architecture. Current Biology 14(19):1739–1746

Via S, Lande R (1985) Genotype-environment interaction and the evolution of phenotypic plasticity. Evolution 39(3):505–522

Vo Van-Zivkovic N, Dinglasan E, Tong J, et al (2025) A large-scale multi-environment study dissecting adult-plant resistance haplotypes for stripe rust resistance in Australian wheat breeding populations. Theoretical and Applied Genetics 138(4):72

Voss-Fels KP, Stahl A, Wittkop B, et al (2019) Breeding improves wheat productivity under contrasting agrochemical input levels. Nature plants 5(7):706–714

Wickham H (2016) Data analysis. In: ggplot2: elegant graphics for data analysis. Springer, p 189–201

Xu S, Hu E, Cai Y, et al (2024) Using clusterProfiler to characterize multiomics data. Nature protocols 19(11):3292–3320

